# A randomized parallel algorithm for efficiently finding near-optimal universal hitting sets

**DOI:** 10.1101/2020.01.17.910513

**Authors:** Barış Ekim, Bonnie Berger, Yaron Orenstein

**Affiliations:** Computer Science and Artificial Intelligence Laboratory, Massachusetts Institute of Technology, Cambridge MA 02139, USA; Department of Mathematics, Massachusetts Institute of Technology, Cambridge MA 02139, USA; School of Electrical and Computer Engineering, Ben-Gurion University of the Negev, Beer-Sheva 8410501, Israel

**Keywords:** Universal hitting sets, parallelization, randomization

## Abstract

As the volume of next generation sequencing data increases, an urgent need for algorithms to efficiently process the data arises. *Universal hitting sets* (UHS) were recently introduced as an alternative to the central idea of minimizers in sequence analysis with the hopes that they could more efficiently address common tasks such as computing hash functions for read overlap, sparse suffix arrays, and Bloom filters. A UHS is a set of *k*-mers that hit every sequence of length *L*, and can thus serve as indices to *L*-long sequences. Unfortunately, methods for computing small UHSs are not yet practical for real-world sequencing instances due to their serial and deterministic nature, which leads to long runtimes and high memory demands when handling typical values of *k* (e.g. *k* > 13). To address this bottleneck, we present two algorithmic innovations to significantly decrease runtime while keeping memory usage low: (i) we leverage advanced theoretical and architectural techniques to parallelize and decrease memory usage in calculating *k*-mer hitting numbers; and (ii) we build upon techniques from randomized Set Cover to select universal *k*-mers much faster. We implemented these innovations in PASHA, the first randomized parallel algorithm for generating near-optimal UHSs, which newly handles *k* > 13. We demonstrate empirically that PASHA produces sets only slightly larger than those of serial deterministic algorithms; moreover, the set size is provably guaranteed to be within a small factor of the optimal size. PASHA’s runtime and memory-usage improvements are orders of magnitude faster than the current best algorithms. We expect our newly-practical construction of UHSs to be adopted in many high-throughput sequence analysis pipelines.

## 1 Introduction

The NIH Sequence Read Archive [8] currently contains over 26 petabases of sequence data. Increased use of sequence-based assays in research and clinical settings creates high computational processing burden; metagenomics studies generate even larger sequencing datasets [19, 17]. New computational ideas are essential to manage and analyze these data. To this end, researchers have turned to *k*-mer-based approaches to more efficiently index datasets [7].

Minimizer techniques were introduced to select *k*-mers from a sequence to allow efficient binning of sequences such that some information about the sequence’s identity is preserved [18]. Formally, given a sequence of length *L* and an integer *k*, its *minimizer* is the lexicographically smallest *k*-mer in it. The method has two key advantages: selected *k*-mers are close; and similar *k*-mers are selected from similar sequences. Minimizers were adopted for biological sequence analysis to design more efficient algorithms, both in terms of memory usage and runtime, by reducing the amount of information processed, while not losing much or any information [12]. The minimizer method has been applied in a large number of settings [4, 20, 6].

Orenstein and Pellow *et al*. [14, 15] generalized and improved upon the minimizer idea by introducing the notion of a *universal hitting set* (UHS). For integers *k* and *L*, set *U*_*k,L*_ is called a universal hitting set of *k*-mers if every possible sequence of length *L* contains at least one *k*-mer from *U*_*k,L*_. Note that a UHS for any given *k* and *L* only needs to be computed once. Their heuristic DOCKS finds a small UHS in two steps: (i) remove a minimum-size set of vertices from a complete de Bruijn graph of order *k* to make it acyclic; and (ii) remove additional vertices to eliminate all (*L* − *k*)-long paths. The removed vertices comprise the UHS. The first step was solved optimally, while the second required a heuristic. The method is limited by runtime to *k* ≤ 13, and thus applicable to only a small subset of minimizer scenarios. Recently, Marçais *et al*. [10] showed that there exists an algorithm to compute a set of *k*-mers that covers every path of length *L* in a de Bruijn graph of order *k*. This algorithm gives an asymptotically optimal solution for a value of *k* approaching *L*. Yet this condition is rarely the case for real applications where 10 ≤ *k* ≤ 30 and 100 ≤ *L* ≤ 300. The results of Marçais *et al.* show that for *k* ≤ 30, the results are far from optimal for fixed *L*. A more recent method by DeBlasio *et al.* [3] can handle larger values of *k*, but with *L* ≤ 21, which is impractical for real applications. Thus, it is still desirable to devise faster algorithms to generate small UHSs.

Here, we present PASHA (Parallel Algorithm for Small Hitting set Approximation), the first randomized parallel algorithm to efficiently generate near-optimal UHSs. Our novel algorithmic contributions are twofold. First, we improve upon the process of calculating vertex hitting numbers, i.e. the number of (*L* − *k*)-long paths they go through. Second, we build upon a randomized parallel algorithm for Set Cover to substantially speedup removal of *k*-mers for the UHS—the major time-limiting step—with a guaranteed approximation ratio on the *k*-mer set size. PASHA performs substantially better than current algorithms at finding a UHS in terms of runtime, with only a small increase in set size; it is consequently applicable to much larger values of *k*. Software and computed sets are available at pasha.csail.mit.edu and github.com/ekimb/pasha.

## 2 Background and Preliminaries

### Preliminary definitions

For *k* ≥ 1 and finite alphabet *Σ*, directed graph *B*_*k*_ = (*V, E*) is a **de Bruijn graph** of order *k* if *V* and *E* represent *k*- and (*k* + 1)-long strings over *Σ*, respectively. An edge may exist from vertex *u* to vertex *v* if the (*k* − 1)-suffix of *u* is the (*k* − 1)-prefix of *v*. For any edge (*u, v*) ∈ *E* with label ℒ, labels of vertices *u* and *v* are the prefix and suffix of length *k* of ℒ, respectively. If a de Bruijn graph contains all possible edges, it is *complete*, and the set of edges represents all possible (*k* + 1)-mers. An *ℓ* = (*L* − *k*)-long path in the graph, i.e. a path of *ℓ* edges, represents an *L*-long sequence over *Σ* (for further details, see [1]).

For any *L*-long string *s* over *Σ, k*-mers set *M* **hits** *s* if there exists a *k*-mer in *M* that is a contiguous substring in *s*. Consequently, **universal hitting set** (UHS) *U*_*k,L*_ is a set of *k*-mers that hits any *L*-long string over *Σ*. A trivial UHS is the set of all *k*-mers, but due to its size (|*Σ*|^*k*^), it does not reduce the computational expense for practical use. Note that a UHS for any given *k* and *L* does not depend on a dataset, but rather needs to be computed only once.

Although the problem of computing a universal hitting set has no known hardness results, there are several NP-hard problems related to it. In particular, the problem of computing a universal hitting set is highly similar, although not identical, to the (*k, L*)-*hitting set* problem, which is the problem of finding a minimum-size *k*-mer set that hits an input set of *L*-long sequences. Orenstein and Pellow *et al*. [14, 15] proved that the (*k, L*)-*hitting set* problem is NP-hard, and consequently developed the near-optimal DOCKS heuristic. DOCKS relies on the Set Cover problem, which is the problem of finding a minimum-size collection of subsets *S*_1_, …, *S*_*k*_ of finite set *U* whose union is *U*.

### The DOCKS heuristic

DOCKS first removes from a complete de Bruijn graph of order *k* a *decycling set*, turning the graph into a directed acyclic graph (DAG). This set of vertices represent a set of *k*-mers that hits all sequences of infinite length. A minimum-size decycling set can be found by Mykkelveit’s algorithm [13] in *O*(|*Σ*|^*k*^) time. Even after all cycles, which represent sequences of infinite length, are removed from the graph, there may still be paths representing sequences of length *L*, which also need to be hit by the UHS. DOCKS removes an additional set of *k*-mers that hits all remaining sequences of length *L*, so that no path representing an *L*-long sequence, i.e. a path of length *ℓ* = *L* − *k*, remains in the graph.

However, finding a minimum-size set of vertices to cover all paths of length *ℓ* in a directed acyclic graph (DAG) is NP-hard [16]. In order to find a small, but not necessarily minimum-size, set of vertices to cover all *ℓ*-long paths, Orenstein and Pellow *et al*. [14, 15] introduced the notion of a *hitting number*, the number of *ℓ*-long paths containing vertex *v*, denoted by *T* (*v, ℓ*). DOCKS uses the hitting number to prioritize removal of vertices that are likely to cover a large number of paths in the graph. This, in fact, is an application of the greedy method for the Set Cover problem, thus guaranteeing an approximation ratio of *O*(1 + log(max_*v*_ *T* (*v, ℓ*))) on the removal of additional *k*-mers.

The hitting numbers for all vertices can be computed efficiently by dynamic programming: For any vertex *v* and 0 ≤ *i* ≤ *ℓ*, DOCKS calculates the number of *i*-long paths starting at *v, D*(*v, i*), and the number of *i*-long paths ending at *v, F* (*v, i*). Then, the hitting number is directly computable by

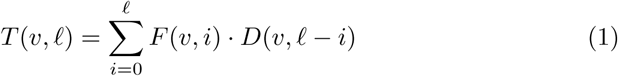

and the dynamic programming calculation in graph *G* = (*V′, E*′) is given by

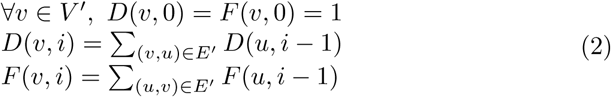

Overall, DOCKS performs two main steps: First, it finds and removes a minimum-size decycling set, turning the graph into a DAG. Then, it iteratively removes vertex *v* with the largest hitting number *T* (*v, ℓ*) until there are no *ℓ*-long paths in the graph. DOCKS is sequential: In each iteration, one vertex with the largest hitting number is removed and added to the UHS output, and the hitting numbers are recalculated. Since the first phase of DOCKS is solved optimally in polynomial time, the bottleneck of the heuristic lies in the removal of the remaining set of *k*-mers to cover all paths of length *ℓ* = *L* − *k* in the graph, which represent all remaining sequences of length *L*.

As an additional heuristic, Orenstein and Pellow *et al*. [14, 15] developed DOCKSany with a similar structure as DOCKS, but instead of removing the vertex that hits the most (*L* − *k*)-long paths, it removes a vertex that hits the most paths in each iteration. This reduces the runtime by a factor of *L*, as calculating the hitting number *T* (*v*) for each vertex can be done in linear time with respect to the size of the graph. DOCKSanyX extends DOCKSany by removing *X* vertices with the largest hitting numbers in each iteration. DOCKSany and DOCKSanyX run faster compared to DOCKS, but the resulting hitting sets are larger.

## 3 Methods

### Overview of the algorithm

Similar to DOCKS, PASHA is run in two phases: First, a minimum-size decycling set is found and removed; then, an additional set of *k*-mers that hits remaining *L*-long sequences is removed. The removal of the decycling set is identical to that of DOCKS; however, in PASHA we introduce randomization and parallelization to efficiently remove the additional set of *k*-mers. We present two novel contributions to efficiently parallelize and randomize the second phase of DOCKS. The first contribution leads to a faster calculation of hitting numbers, thus reducing the runtime of each iteration. The second contribution leads to selecting multiple vertices for removal at each iteration, thus reducing the number of iterations to obtain a graph with no (*L* − *k*)-long paths. Together, the two contributions provide orthogonal improvements in runtime.

### Improved hitting number calculation

#### Memory usage improvements

We reduce memory usage through algorithmic and technical advances. Instead of storing the number of *i*-long paths for 0 ≤ *i* ≤ *ℓ* in both *F* and *D*, we apply the following approach (Algorithm 1): We compute *D* for all *v* ∈ *V* and 0 ≤ *i* ≤ *ℓ*. Then, while computing the hitting number, we calculate *F* for iteration *i*. For this aim, we define two arrays: *F*_*curr*_ and *F*_*prev*_, to store only two instances of *i*-long path counts for each vertex: The current and previous iterations. Then, for some *j*, we compute *F*_*curr*_ based on *F*_*prev*_, set *F*_*prev*_ = *F*_*curr*_, and add *F*_*curr*_(*v*) · *D*(*v, ℓ* − *j*) to the hitting number sum. Lastly, we increase *j*, and repeat the procedure, adding the computed hitting numbers iteratively. This approach allows the reduction of matrix *F*, since in each iteration we are storing only two arrays, *F*_*curr*_ and *F*_*prev*_, instead of the original *F* matrix consisting of *ℓ* + 1 arrays. Therefore, we are able to reduce memory usage by close to half, with no change in runtime.

To further reduce memory usage, we use `float` variable type (of size 4 bytes) instead of `double` variable type (of size 8 bytes). The number of paths kept in *F* and *D* increase exponentially with *i*, the length of the paths. To be able to use the 8 bit exponent field, we initialize *F* and *D* to `float` minimum positive value. This does not disturb algorithm correctness, as path counting is only scaled to some arbitrary unit value, which may be 2^−149^, the smallest positive value that can be represented by `float`. This is done in order to account for the high numbers that path counts can reach. The remaining main memory bottleneck is matrix *D*, whose size is 4 · 4^*k*^ · (*ℓ* + 1) bytes.

Lastly, we utilized the property of a complete de Bruijn graph of order *k* being the line graph of a de Bruijn graph of order *k* − 1. While all *k*-mers are represented as the set of vertices in the graph of order *k*, they are represented as edges in the graph of order *k* − 1. If we remove edges of a de Bruijn graph of order *k* − 1, instead of vertices in a graph of order *k*, we can reduce memory usage by another factor of |*Σ*|. In our implementation we compute *D* and *F* for all vertices of a graph of order *k* − 1, and calculate hitting numbers for edges. Thus, the bottleneck of the memory usage is reduced to 4 · 4^*k*−1^ · (*ℓ* + 1) bytes.

#### Runtime reduction by parallelization

We parallelize the calculation of the hitting numbers to achieve a constant factor reduction in runtime. The calculation of *i*-long paths through vertex *v* only depends on the previously calculated matrices for the (*i* − 1)-long paths through all vertices adjacent to *v* (Equation 2). Therefore, for some *i*, we can compute *D*(*v, i*) and *F* (*v, i*) for all vertices in *V′* in parallel, where *V′* is the set of vertices left after the removal of the decycling set. In addition, we can calculate the hitting number *T* (*v, ℓ*) for all vertices *V′* in parallel (similar to computing *D* and *F*), since the calculation does not depend on the hitting number of any other vertex (we call this parallel variant PDOCKS for the purpose of comparison with PASHA). We note that for DOCKSany and DOCKSanyX, the calculations of hitting numbers for each vertex cannot be computed in parallel, since the number of paths starting and ending at each vertex both depend on those of the previous vertex in topological order.

##### Algorithm 1 Improved hitting numbers calculation. *Input: G* = (*V, E*)

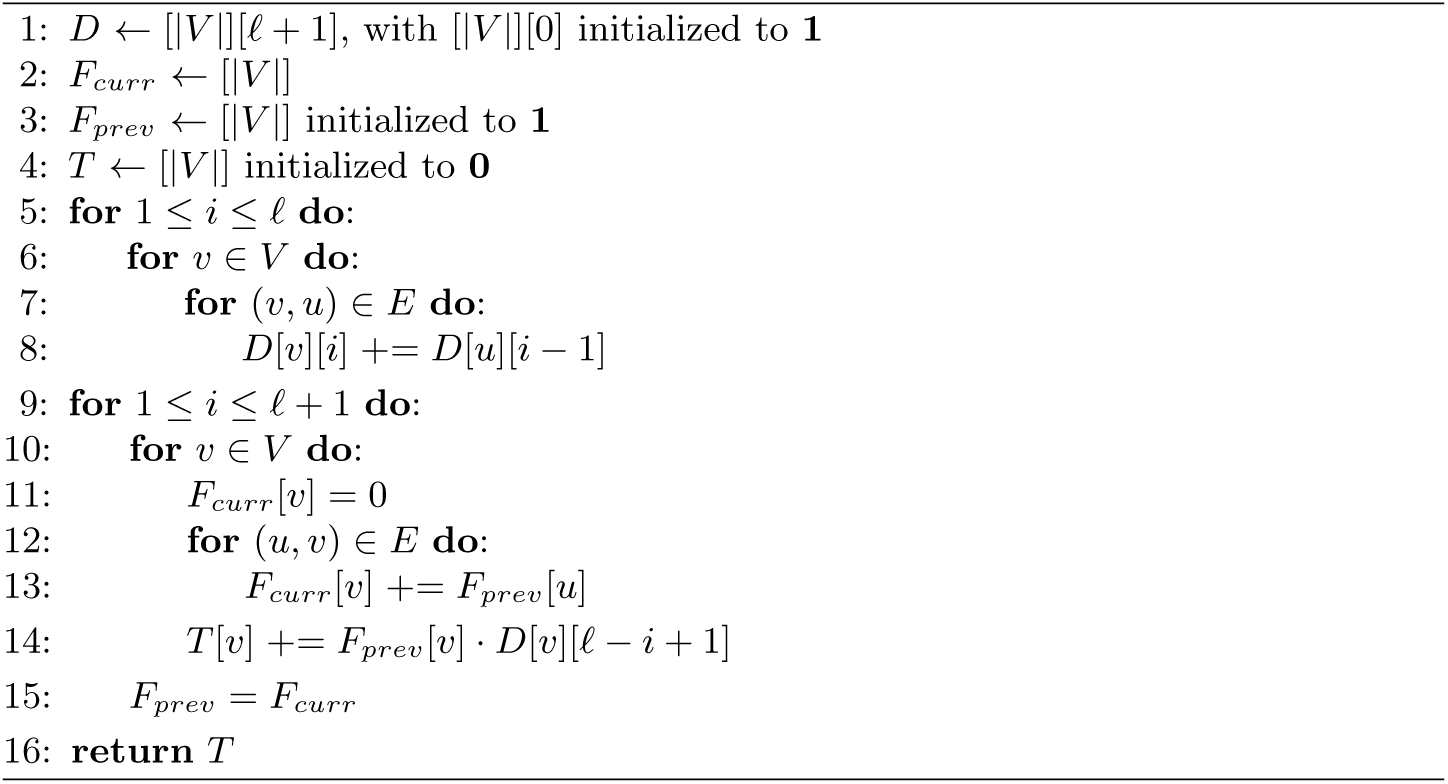

### Parallel randomized *k*-mer selection

Our goal is to find a minimum-size set of vertices that covers all *ℓ*-long paths. We can represent the remaining graph as an instance of the Set Cover problem. While the greedy algorithm for the second phase of DOCKS is serial, we will show that we can devise a parallel algorithm, which is close to the greedy algorithm in terms of performance guarantees, by picking a large set of vertices that cover nearly as many paths as the vertices that the greedy algorithm picks one by one.

In PASHA, instead of removing the vertex with the maximum hitting number in each iteration, we consider a set of vertices for removal with hitting numbers within an interval, and pick vertices in this set independently with constant probability. Considering vertices within an interval allows us to efficiently introduce randomization while still emulating the deterministic algorithm. Picking vertices independently in each iteration enables parallelization of the procedure. Our randomized parallel algorithm for the second phase of the UHS problem adapts that of Berger *et al*. [2] for the original Set Cover problem.

#### The UHS selection procedure

The input includes graph *G* = (*V, E*) and randomization variables 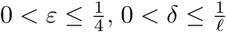 (Algorithm 2). Let function calcHit() calculate the hitting numbers for all vertices, and return the maximum hitting number (line 2). We set *t* = ⌈log_1+*ε*_ *T*_*max*_⌉ (line 3), and run a series of steps from *t*, iteratively decreasing *t* by 1. In step *t*, we first calculate the hitting numbers of all vertices (line 5); then, we define vertex set *S* to contain vertices with a hitting number between (1 + *ε*)^*t*−1^ and (1 + *ε*)^*t*^ for potential removal (lines 8-9).

Let *P*_*S*_ be the sum of all hitting numbers of the vertices in *S*, i.e. *P*_*S*_ = ∑_*v*∈*S*_ *T* (*v, ℓ*) (line 10). In each step, if the hitting number for vertex *v* is more than a *δ*^3^ fraction of *P*_*S*_, i.e. *T* (*v, ℓ*) ≥ *δ*^3^*P*_*S*_, we add *v* to the picked vertex set *V*_*t*_ (lines 11-13). For vertices with a hitting number smaller than *δ*^3^*P*_*S*_, we pairwise independently pick them with probability 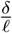. We test the vertices in pairs to impose pairwise independence: If an unpicked vertex *u* satisfies the probability 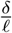, we choose another unpicked vertex *v* and test the same probability 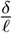. If both are satisfied, we add both vertices to the picked vertex set *V*_*t*_; if not, neither of them are added to the set (lines 14-16). This serves as a bound on the probability of picking a vertex. If the sum of hitting numbers of the vertices in set *V*_*t*_ is at least |*V*_*t*_| (1 + *ε*)^*t*^(1−4*δ*−2*ε*), we add the vertices to the output set, remove them from the graph, and decrease *t* by 1 (lines 17-20). The next iteration runs with decreased *t*. Otherwise, we rerun the selection procedure without decreasing *t*.

##### Algorithm 2 The selection procedure. *Input*: 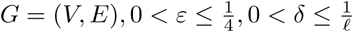

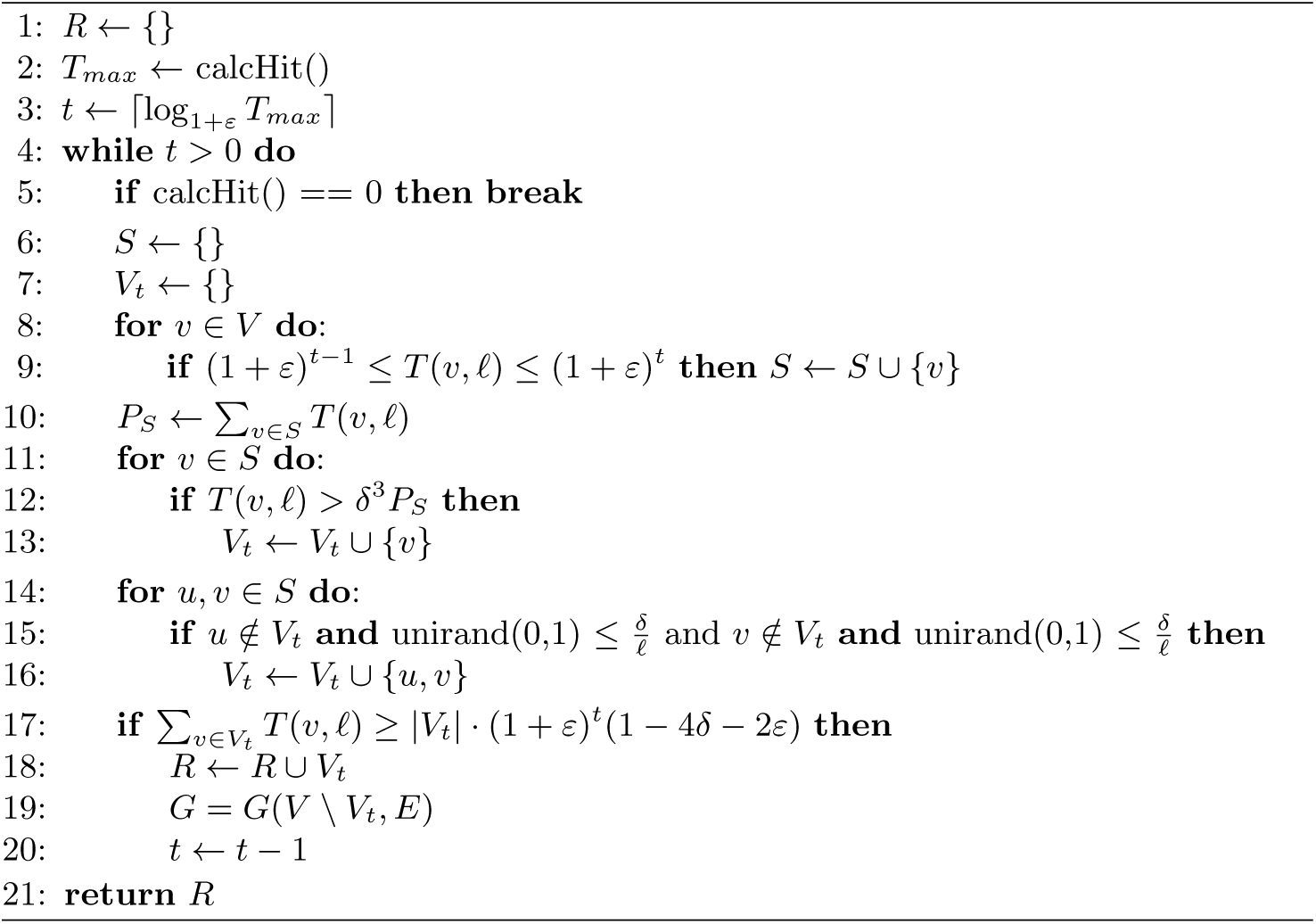

#### Performance guarantees

At step *t*, we add the selected vertex set *V*_*t*_ to the output set if 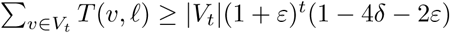. Otherwise, we rerun the selection procedure with the same value of *t*. We show in Appendix A that with high probability, 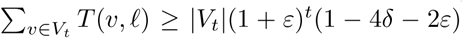. We also show that PASHA produces a cover *α*(1 + log *T*_*max*_) times the optimal size, where *α* = 1*/*(1−4*δ*−2*ε*). In Appendix B, we give the asymptotic number of the selection steps and prove the average runtime complexity of the algorithm. Performance summaries in terms of theoretical runtime and approximation ratio are in Table 1.

**Table 1.**
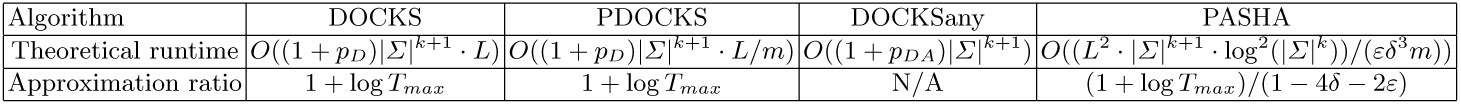
Summary of theoretical results for the second phase of different algorithms for generating a set of *k*-mers hitting all *L*-long sequences. PDOCKS is DOCKS with the improved hitting number calculation, i.e. greedy removal of one vertex at each iteration. *p*_*D*_, *p*_*DA*_ denote the total number of picked vertices for DOCKS/PDOCKS and DOCKSany, respectively. *m* denotes the number of parallel threads used, *T*_*max*_ the maximum vertex hitting number, and *ϵ* and *δ* PASHA’s randomization parameters.

## 4 Results

### PASHA outperforms extant algorithms for *k ≤* 13

We compared PASHA and PDOCKS to extant methods on several combinations of *k* and *L*. We ran DOCKS, DOCKSany, PDOCKS, and PASHA over 5 ≤ *k* ≤ 10, DOCKSanyX, PDOCKS, and PASHA for *k* = 11 and *X* = 10, and PASHA and DOCKSanyX for *X* = 100, 1000 for *k* = 12, 13 respectively, for 20 ≤ *L* ≤ 200. We say that an algorithm is *limited by runtime* if for some value of *k* ≤ 13 and for *L* = 100, its runtime exceeds 1 day (86400 seconds), in which case we stopped the operation and excluded the method from the results for the corresponding value of *k*. While running PASHA, we set *δ* = 1*/ℓ*, and 1−4*δ*−2*ε* = 1*/*2 to set an emulation ratio *α* = 2 (see Section 3 and Appendix A). The methods were benchmarked on a 24-CPU Intel Xeon Gold (2.10GHz) with 754GB of RAM. We ran all tests using all available cores (*m* = 24 in Table 1).

#### Comparing runtimes and UHS sizes

We ran DOCKS, PDOCKS, DOCKSany, and PASHA for *k* = 10 and 20 ≤ *L* ≤ 200. As seen in Figure 1A, DOCKS has a significantly higher runtime than the parallel variant PDOCKS, while producing identical sets (Figure 1B). For small values of *L*, DOCKSany produces the largest UHSs compared to other methods, and as *L* increases, the differences in both runtime and UHS size for all methods decrease, since there are fewer *k*-mers to add to the removed decycling set to produce a UHS.

**Fig. 1.**
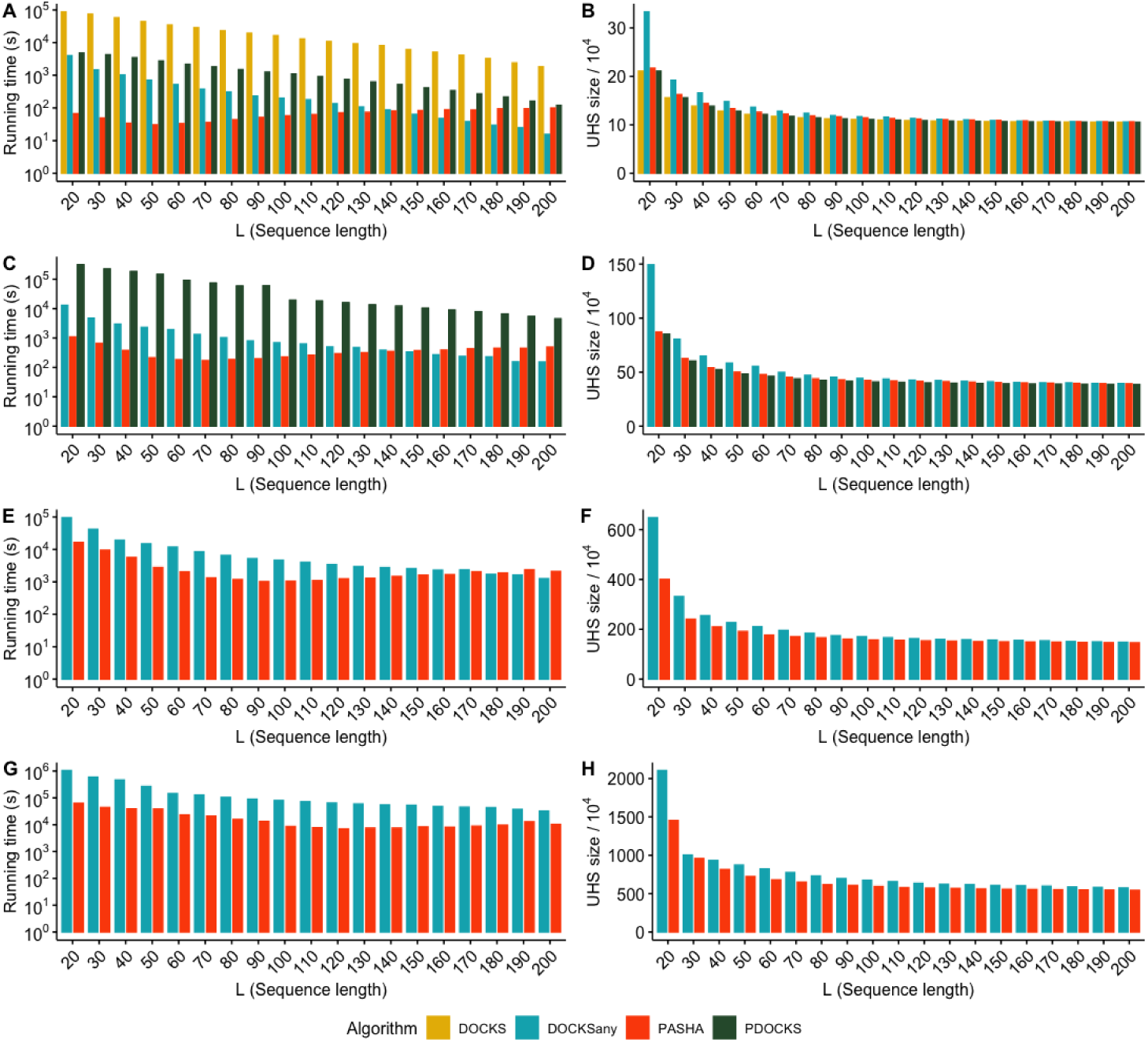
Runtimes (left) and UHS sizes (divided by 10^4^, right) for values of *k* = 10 (A, B), 11 (C, D), 12 (E, F), and 13 (G, H) and 20 ≤ *L* ≤ 200 for the different methods. Note that the y-axes for runtimes are in logarithmic scale.

We ran PDOCKS, DOCKSany10, and PASHA for *k* = 11 and 20 ≤ *L* ≤ 200. As seen in Figure 1C, for small values of *L*, both PDOCKS and DOCKSany10 have significantly higher runtimes than PASHA; while for larger *L*, DOCKSany10 and PASHA are comparable in their runtimes (with PASHA being negligibly slower). In Figure 1D, we observe that PDOCKS computes the smallest sets for all values of *L*. Indeed, its guaranteed approximation ratio is the smallest among all three benchmarked methods. While the set sizes for all methods converge to the same value for larger *L*, DOCKSany10 produces the largest UHSs for small values of *L*, in which case PASHA and PDOCKS are preferable.

PASHA’s runtime behaves differently than that of other methods. For all methods but PASHA, runtime decreases as *L* increases. Instead of gradually decreasing with *L*, PASHA’s runtime gradually decreases up to *L* = 70, at which it starts to increase at a much slower rate. This is explained by the asymptotic complexity of PASHA (Table 1). Since computing a UHS for small *L* requires a larger number of vertices to be removed, the decrease in runtime with increasing *L* up to *L* = 70 is significant; however, due to PASHA’s asymptotic complexity being quadratic with respect to *L*, we see a small increase from *L* = 70 to *L* = 200. All other methods depend linearly on the number of removed vertices, which decreases as *L* increases.

Despite the significant decrease in runtime in PDOCKS compared to DOCKS, PDOCKS was still limited by runtime to *k* ≤ 12. Therefore, we ran DOCKSany100 and PASHA for *k* = 12 and 20 ≤ *L* ≤ 200. As seen in Figures 1E and 1F, both methods follow a similar trend as in *k* = 11, with DOCKSany100 being significantly slower and generating significantly larger UHSs for small values of *L*. For larger values of *L*, DOCKSany100 is slightly faster, while PASHA produces sets that are slightly smaller.

At *k* = 13 we observed the superior performance of PASHA over DOCKSany1000 in both runtime and set size for all values of *L*. We ran DOCKSany1000 and PASHA for *k* = 13 and 20 ≤ *L* ≤ 200. As seen in Figures 1G and 1H, DOCKSany1000 produces larger sets and is significantly slower compared to PASHA for all values of *L*. This result demonstrates that the slow increase in runtime for PASHA compared to other algorithms for *k* < 13 does not have a significant effect on runtime for larger values of *k*.

### PASHA enables UHS for *k* = 14, 15, 16

Since all existing algorithms and PDOCKS are limited by runtime to *k* ≤ 13, we report the first UHSs for 14 ≤ *k* ≤ 16 and *L* = 100 computed using PASHA, run on a 24-CPU Intel Xeon Gold (2.10GHz) with 754GB of RAM using all 24 cores. Figure 2 shows runtimes and sizes of the sets computed by PASHA.

**Fig. 2.**
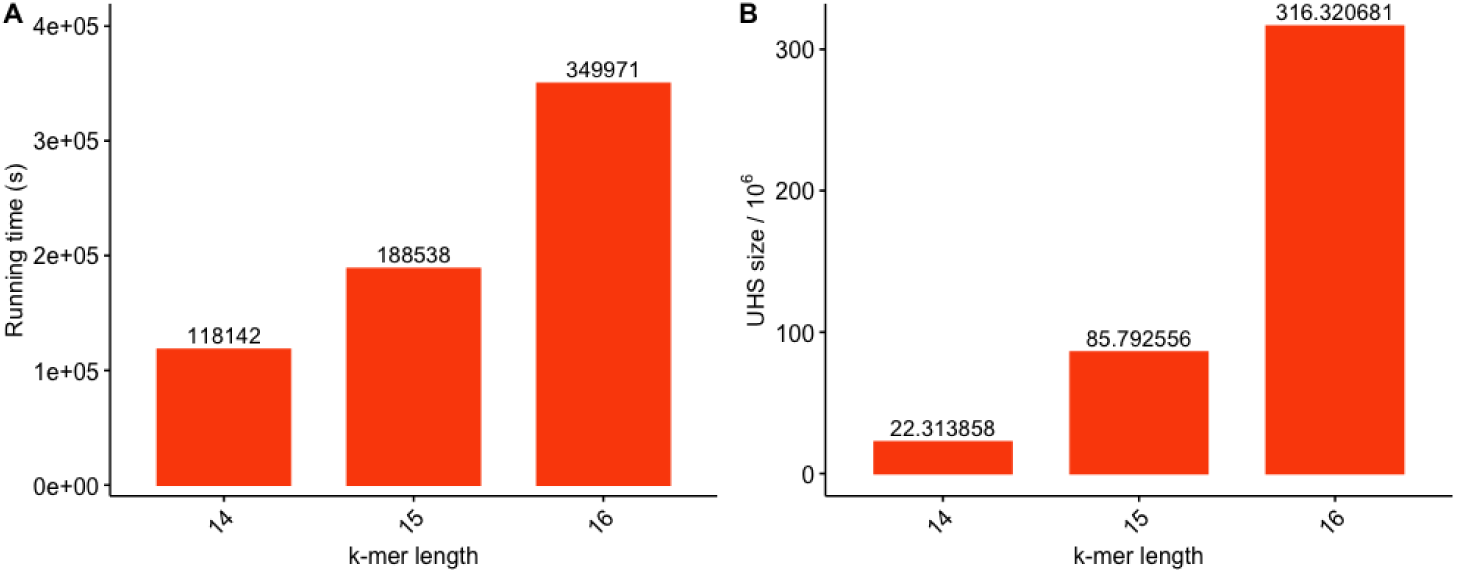
Runtimes (A) and UHS sizes (divided by 10^6^) (B) for 14 ≤ *k* ≤ 16 and *L* = 100 for PASHA. Note that the y-axis for runtime is in logarithmic scale.

### Density comparisons for the different methods

In addition to runtimes and UHS sizes, we report values of another measure of UHS performance known as *density*. The *density* of the minimizers scheme *d*(*M, S, k*) is the fraction of selected *k*-mers’ positions over the number of *k*-mers in the sequence. Formally, the density of scheme *M* over sequence *S* is defined as

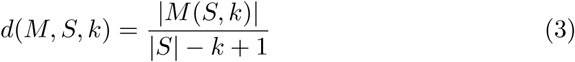

where *M* (*S, k*) is the set of positions of the *k*-mers selected over sequence *S*.

We calculate densities for a UHS by selecting the lexicographically smallest *k*-mer that is in the UHS within each window of *L* − *k* + 1 consecutive *k*-mers, since at least one *k*-mer is guaranteed to be in each such window. Marçais *et al.* [11] showed that using UHSs for *k*-mer selection in this manner yields smaller densities than lexicographic or random minimizer selection schemes. Therefore, we do not report comparisons between UHSs and minimizer schemes, but rather comparisons among UHSs constructed by different methods.

Marçais *et al.* [11] also showed that the expected density of a minimizers scheme for any *k* and window size *L* − *k* + 1 is equal to the density of the minimizers scheme on a de Bruijn sequence of order *L*. This allows for exact calculation of expected density for any *k*-mer selection procedure. However, for 14 ≤ *k* ≤ 16 we calculated UHSs only for *L* = 100, and iterating over a de Bruijn sequence of order 100 is infeasible. Therefore, we computed the approximate expected density on long random sequences, since the computed expected density on these sequences converges to the expected density [11]. In addition, we computed the density of different methods on the entire human reference genome (GRCh38).

We computed the density values of UHSs generated by PDOCKS, DOCKSany, and PASHA over 10 random sequences of length 10^6^, and the entire human reference genome (GRCh38), for 5 ≤ *k* ≤ 16 and *L* = 100, when a UHS was available for such (*k, L*) combination.

As seen in Figure 3, the differences in both approximate expected density and density computed on the human reference genome are negligible when comparing UHSs generated by the different methods. For most values of *k*, DOCKS yields the smallest approximate expected density and human genome density values, while DOCKSany generally yields lower human genome density values, but higher expected density values than PASHA. For *k* ≤ 6, the UHS is only the decycling set; therefore, density values for these values of *k* are identical for the different methods.

**Fig. 3.**
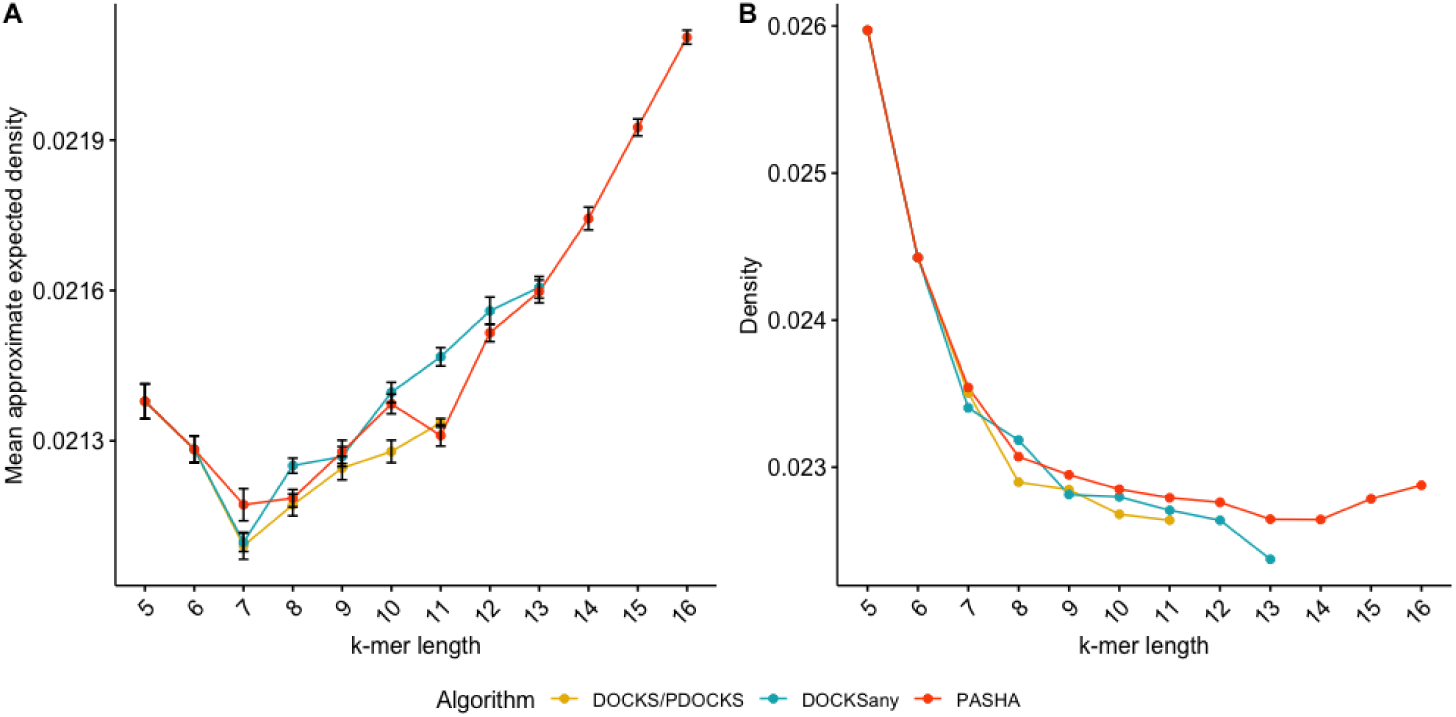
Mean approximate expected density (A), and density on the human reference genome (B) for different methods, for 5 ≤ *k* ≤ 16 and *L* = 100. Error bars represent one standard deviation from the mean across 10 random sequences of length 10^6^. Density is the fraction of selected *k*-mer positions over the number of *k*-mers in the sequence.

Since there is no significant difference in the density of the UHSs generated by the different methods, other criteria, such as runtime and set size, are relevant when evaluating the performance of the methods: As *k* increases, PASHA produces sets that are only slightly smaller or larger in density, but significantly smaller in size and significantly faster than extant methods.

## 5 Discussion

We presented an efficient randomized parallel algorithm for generating a small set of *k*-mers that hits every possible sequence of length *L* and produces a set that is a small guaranteed factor away from the optimal set size. Since the runtime of DOCKS variants and PASHA depend exponentially on *k*, these greedy heuristics are eventually limited by runtime. However, using these heuristics in conjunction with parallelization, we are newly able to compute UHSs for values of *k* and *L* large enough for most biological applications.

The improvements in runtime for the hitting number calculation are due to parallelization of the dynamic programming phase, which is the bottleneck in sequential DOCKS variants. A minimum-size set that hits all infinite-length sequences is optimally and rapidly removed; however, the remaining sequences of length *L* are calculated and removed in time polynomial in the output size. We show that a constant factor reduction is beneficial in mitigating this bottleneck for practical use. In addition, we reduce the memory usage of this phase by theoretical and technical advancements. Last, we build on a randomized parallel algorithm for Set Cover to significantly speed up vertex selection. The randomized algorithm can be derandomized, while preserving the same approximation ratio, since it requires only pairwise independence of the random variables [2].

One main open problem still remains from this work. Although the randomized approximation algorithm enables us to generate a UHS more efficiently, the hitting numbers still need to be calculated at each iteration. The task of computing hitting numbers remains as the bottleneck in computing a UHS. Is there a more efficient way of calculating hitting numbers than the dynamic programming calculation done in DOCKS and PASHA?

As for long reads, which are becoming more popular for genome assembly tasks, a *k*-mer set that hits all infinite long sequences, as computed optimally by Mykkelveit’s algorithm [13], is enough due to the length of these long read sequences. Still, due to the inaccuracies and high cost of long read sequencing compared to short read sequencing, the latter is still the prevailing method to produce sequencing data, and is expected to remain so for the near future.

We expect the efficient calculation of UHSs to lead to improvements in sequence analysis and construction of space-efficient data structures. Unfortunately, previous methods were limited to small values of *k*, thus allowing application to only a small subset of sequence analysis tasks. As there is an inherent exponential dependency on *k* in terms of both runtime and memory, efficiency in calculating these sets is crucial. We expect that the UHSs newly-enabled by PASHA for *k* > 13 will be useful in improving various applications in genomics.

## 6 Conclusion

We developed a novel randomized parallel algorithm PASHA to compute a small set of *k*-mers which together hit every sequence of length *L*. It is based on two algorithmic innovations: (i) improved calculation of hitting numbers through paralleization and memory reduction; and (ii) randomized parallel selection of additional *k*-mers to remove. We demonstrated the scalability of PASHA to larger values of *k* up to 16. Notably, the universal hitting sets need to be computed only once, and can then be used in many sequence analysis applications. We expect our algorithms to be an essential part of the sequence analysis toolkit.

## 7 Acknowledgments

This work was supported by NIH grant R01GM081871 to B.B. B.E. was supported by the MISTI MIT-Israel program at MIT and Ben-Gurion University of the Negev. We gratefully acknowledge the support of Intel Corporation for giving access to the Intel® AI DevCloud platform used for part of this work.

## A Emulating the greedy algorithm

The greedy Set Cover algorithm was developed independently by Johnson and Lovász for unweighted vertices [5, 9]. Lovász [9] proved:

### Theorem 1.

*The greedy algorithm for Set Cover outputs cover R with* |*R*| ≤ (1 + log *T*_*max*_)|*OPT* |, *where T*_*max*_ *is the maximum cardinality set.*

We adapt a definition for an algorithm emulating the greedy algorithm for the Set Cover problem to the second phase of DOCKS [2]. We say that an algorithm for the second phase of DOCKS *α***-emulates** the greedy algorithm if it outputs a set of vertices serially, during which it selects vertex set *A* such that

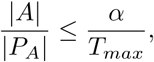

where *P*_*A*_ is the set of *ℓ*-long paths covered by *A*. Using this definition, we come up with a near-optimal approximation by the following theorem:

### Theorem 2.

*An algorithm for the second phase of DOCKS that α-emulates the greedy algorithm produces cover R* ⊆ *V with* |*R*| ≤ *α*(1 + log *T*_*max*_) |*OPT*|, *where OPT is the optimal cover.*

*Proof.* We define the *cost* of covering path *p* as 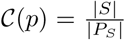, where *S* is the set of vertices selected in the selection step in which *p* was covered, and *P*_*S*_ the set of *ℓ*-long paths covered by *S*. Then, 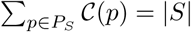.

Let *P*_*ℓ*_ be set of all *ℓ*-long paths in *G*. A **fractional cover** of graph *G* = (*V, E*) is function ℱ : *V* → {0, 1} s.t. for all *p* ∈ *P*_*ℓ*_, ∑_*v*∈*p*_ ℱ (*v*) ≥ 1. The optimal cover ℱ _*OPT*_ has minimum ∑_*v*∈*V*_ ℱ _*OPT*_ (*v*).

Let ℱ be such an optimal fractional cover. The size of the cover produced is

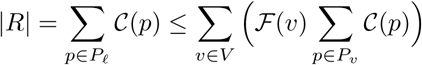

where *P*_*v*_ is the set of all *ℓ*-long paths through vertex *v*.

### Lemma 1.

*There are at most* 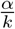 *paths p* ∈ *P*_*v*_ *such that* 𝒞(*p*) ≥ *k for any v, k.*

*Proof.* Assume the contrary: Before such path *p* is covered, 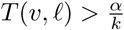. Thus,

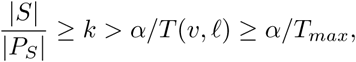

contradicting the definition.

Suppose we rank the *T* (*v, ℓ*) paths *p* ∈ *P*_*v*_ by decreasing order of 𝒞(*p*). From the above remark, if the *i*th path has cost *k*, then *i* ≤ *α/k*. Then, we can write

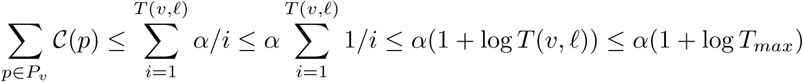

Then,

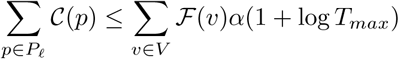

and finally

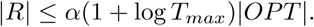

In PASHA, we ensure that in step *t*, the sum of vertex hitting numbers of selected vertex set *V*_*t*_ is at least | *V*_*t*_| (1 + *ε*)^*t*^(1 −4*δ* −2*ε*). We now show that this is satisfied with high probability in each step.

### Theorem 3.

*With probability at least 1/2, the sum of vertex hitting numbers of selected vertex set V*_*t*_ *at step t is at least* |*V*_*t*_|(1 + *ε*)^*t*^(1 − 4*δ* − 2*ε*).

*Proof.* For any vertex *v* in selected vertex set *V*_*t*_ at step *t*, let *X*_*v*_ be an indicator variable for the random event that vertex *v* is picked, and 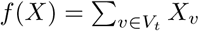.

Note that Var[*f*(*X*)] ≤ |*V*_*t*_|·*δ/ℓ*, and |*V*_*t*_ |> *ℓ/δ*^3^, since we are given that no vertex covers a *δ*^3^ fraction of the *ℓ*-long paths covered by the vertices in *V*_*t*_. By Chebyshev’s inequality, for any *k* ≥ 0,

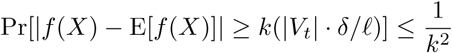

and with probability 3/4,

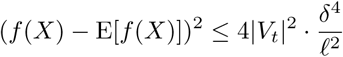

and

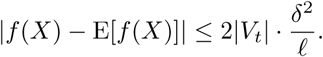

Let 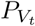 denote the set of *ℓ*-long paths covered by vertex set *V*_*t*_. Then,

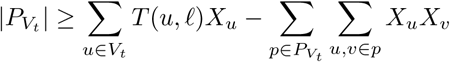

We know that 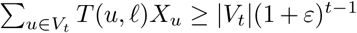. Then,

*T* (*u, ℓ*)*X*_*u*_ ≥ |*V*_*t*_|(1 + *ε*)^*t*−1^, which is bounded below by 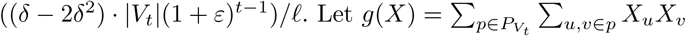. Then,

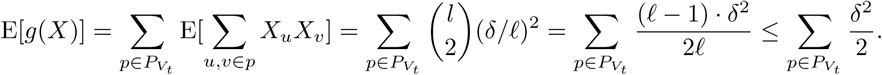

Hence, with probability at least 3/4,

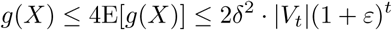

Both events hold with probability at least 1/2, and the sum of vertex hitting numbers is at least

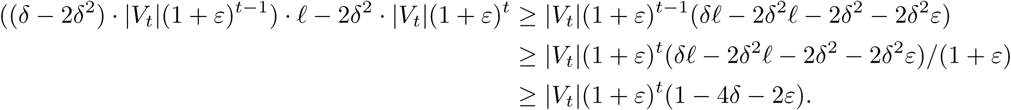

## B Runtime analysis

Here, we show the number of the selection steps and the average-time asymptotic complexity of PASHA.

### Lemma 2.

*The number of selection steps is O*(log |*V* | log |*P*_*ℓ*_ |*/*(*εδ*^3^*m*)).

*Proof.* The number of steps is *O*(log |*V*| */ε*), and within each step, there are *O*(log |*P*_*S*_| */*(*δ*^3^*m*)) selection steps (where *P*_*S*_ is the sum of vertex hitting numbers of the vertex set *S* for that step and *m* the number of threads used), since we are guaranteed to remove a *δ*^3^ fraction of the paths during that step. Overall, there are *O*(log |*V* | log |*P*_*ℓ*_ |*/*(*εδ*^3^*m*)) selection steps.

### Theorem 4.

*For ε* < 1, *there is an approximation algorithm for the second phase of DOCKS that runs in O*((*L*^2^ ·|*Σ*|^*k*+1^ · log^2^(|*Σ*|^*k*^))*/*(*εδ*^3^*m*)) *average time, where m is the number of threads used, and produces a cover of size at most* (1 + *ε*)(1 + log *T*_*max*_) *times the optimal size.*

*Proof.* Follows immediately from Theorem 2 and Lemma 2.

